# A comprehensive CRISPR screen of the Drosophila glutamate receptome reveals Ekar as a selective regulator of presynaptic homeostatic plasticity

**DOI:** 10.64898/2026.03.20.713276

**Authors:** Joshua C. Martinez, Chun Chien, Wanying Dong, Nancy L. Tran, Allison Chang, Hila Zak, Svara Shah, Galit Shohat-Ophir, Dion Dickman

**Affiliations:** Department of Neurobiology, University of Southern California, Los Angeles, CA, USA; The Mina & Everad Goodman Faculty of Life Science, the Multidisciplinary Brain Research Center and the Institute for Nanotechnology and Advanced Materials, Bar-Ilan University, Israel

**Author notes:** Correspondence: Dion Dickman, Department of Neurobiology, University of Southern California, Los Angeles, CA 90089. Faculty of Engineering, Bar-Ilan University, Israel (Hila Zak). These authors contributed equally (Joshua C. Martinez, Chun Chien).

**Keywords:** Glutamate receptor, homeostatic plasticity, neuromuscular junction, Drosophila

## Abstract

Homeostatic mechanisms protect synapses from destabilizing challenges throughout an organism’s lifespan, ensuring stable yet flexible neural network activity. To delineate the molecular basis of presynaptic homeostatic potentiation (PHP), we conducted a comprehensive, *in vivo* CRISPR/Cas9-based screen of all 16 glutamate receptor (GluR) genes encoded in the Drosophila genome. We first generated a complete expression atlas across larval and adult stages, identifying nine GluRs expressed in presynaptic motor neurons. We then generated null mutants for all 16 GluRs and screened them at the larval neuromuscular junction. While the loss of any single presynaptic GluR did not affect baseline synaptic growth or neurotransmission, our screen revealed a selective and critical requirement for the kainate receptor subunit *ekar* in the expression of chronic PHP. Further genetic analysis indicates that Ekar functions coordinately with the kainate receptor subunits KaiRID and Ukar within a shared pathway to promote this plasticity. Mechanistically, Ekar acts downstream of active zone remodeling to drive the homeostatic enhancement of presynaptic Ca^2+^ influx, which is the defining feature of chronic PHP. Together, this genome-wide analysis establishes a definitive functional atlas for the Drosophila glutamate receptome and highlights a specialized, essential role for Ekar in stabilizing long-term synaptic homeostasis.

## INTRODUCTION

Neural circuits must maintain stable function despite the constant perturbations inherent to development, maturation, and disease. This essential stability is actively enforced by conserved homeostatic plasticity mechanisms (Turrigiano 2012; Davis 2006; Davis 2013; Pozo and Goda 2010). A premier example illustrating this adaptation is Presynaptic Homeostatic Potentiation (PHP), wherein the pharmacological or genetic disruption of postsynaptic glutamate receptors (GluRs) triggers a compensatory increase in presynaptic glutamate release to restore baseline synaptic strength (Frank 2014; Goel and Dickman, 2021). Evolutionarily conserved across neuromuscular junctions (NMJs) of flies, rodents, and humans (Cull-Candy et al; 1980, Plomp et al, 1992; Petersen et al, 1997; Orr et al, 2020; Engisch et al, 2022), PHP has also recently been characterized in the mammalian central nervous system (Delvendahl et al, 2019; Chipman et al, 2022; Chipman et al, 2025). The Drosophila NMJ has emerged as a powerful, genetically tractable model system to deconstruct the molecular architecture of this synaptic resilience (Frank and Müller 2020; Davis and Müller 2015; He and Dickman et al, 2025). Yet, despite PHP being a defining example of synaptic homeostasis, the comprehensive presynaptic machinery necessary to execute this potentiation remains incompletely understood. Moreover, recent evidence demonstrates that acute and chronic forms of PHP target distinct motor inputs (Chien et al, 2025; Newman et al, 2017), raising the possibility that dedicated presynaptic signaling systems may underlie this input-specificity.

Previous forward genetic screens using the Drosophila NMJ have identified several key molecules required within the presynaptic motor neuron for PHP expression (Dickman and Davis, 2009; Müller et al, 2011; Dickman et al, 2012; Kikuma et al, 2019; Goel et al, 2019). Notably, these efforts uncovered the presynaptic kainate-type glutamate receptor subunit, KaiRID, which functions as a glutamate autoreceptor, localizing proximal to active zones (Kiragasi et al, 2017). Here, KaiRID was shown to respond to synaptic glutamate to drive increased transmitter release under conditions of limiting Ca^2+^ and during PHP. While *in vitro* studies show that KaiRID can form functional homomeric channels (Li et al, 2016), ionotropic GluRs typically operate as heteromeric assemblies *in vivo*. Indeed, subsequent work identified additional presynaptic GluR machinery required for PHP, including the GluR subunit *ukar*, and the auxiliary GluR subunits dSol-1 and Neto-α (Nguyen et al, 2024; Kiragasi et al, 2020; Han et al, 2020). These auxiliary proteins are critical for homeostatic signaling; dSol-1, for instance, directs the rapid accumulation of KaiRID at release sites during homeostatic signaling (Kiragasi et al, 2020). Collectively, these findings establish that a presynaptic GluR complex tunes both baseline neurotransmission and homeostatic plasticity. Nevertheless, the full complement and molecular composition of GluR subunits orchestrating PHP in motor neurons remains undefined.

To resolve this, we systematically investigated the complete repertoire of GluRs contributing to PHP signaling by executing an *in vivo* CRISPR/Cas9 screen of all 16 Drosophila GluR subunits. Our approach was designed to address three central questions: **1)** Which of the 16 GluRs are expressed at the larval NMJ? **2)** Do these subunits govern synaptic growth or baseline transmission? And **3)** Are additional GluR subunits required for the expression of PHP? We first constructed a comprehensive expression atlas, mapping all 16 GluRs across motor neurons, postsynaptic muscle, other tissues, and developmental stages. We next generated null mutations for each receptor via CRISPR/Cas9 gene editing to assess their functional contributions. While our analysis reveals that most individual GluRs are dispensable for baseline synaptic architecture and function, we identified a critical, previously unknown requirement for the kainate receptor subunit *ekar*. Remarkably, Ekar represents the first GluR subunit selectively required for the “chronic” expression of PHP – the persistent compensatory state driven by the congenital genetic loss of postsynaptic receptors.

## RESULTS

### Expression mapping and basic characterization of all Drosophila GluRs

To determine whether any additional GluR subunits are necessary for PHP expression, we first evaluated the full complement of GluRs encoded in the Drosophila genome (the “GluR receptome”). This phylogenetic analysis confirmed that 16 GluRs are encoded in Drosophila, comprising evolutionarily conserved classes: ten Kainate-type, two AMPA-type, two NMDA-type, one metabotropic, and one glutamate-gated chloride receptor (**Fig. 1A**). To map the tissue-specific expression of each receptor, we obtained or generated promoter-GAL4 fusions (Table S2; Kondo et al, 2020) and examined expression patterns using a membrane-targeted GFP reporter (CD4::tdGFP). Initially probing the central nervous system, we found that 9 of the 16 receptors are expressed in the larval brain, while 11 are expressed in the adult brain (**Fig. S1**).

**Figure 1:**
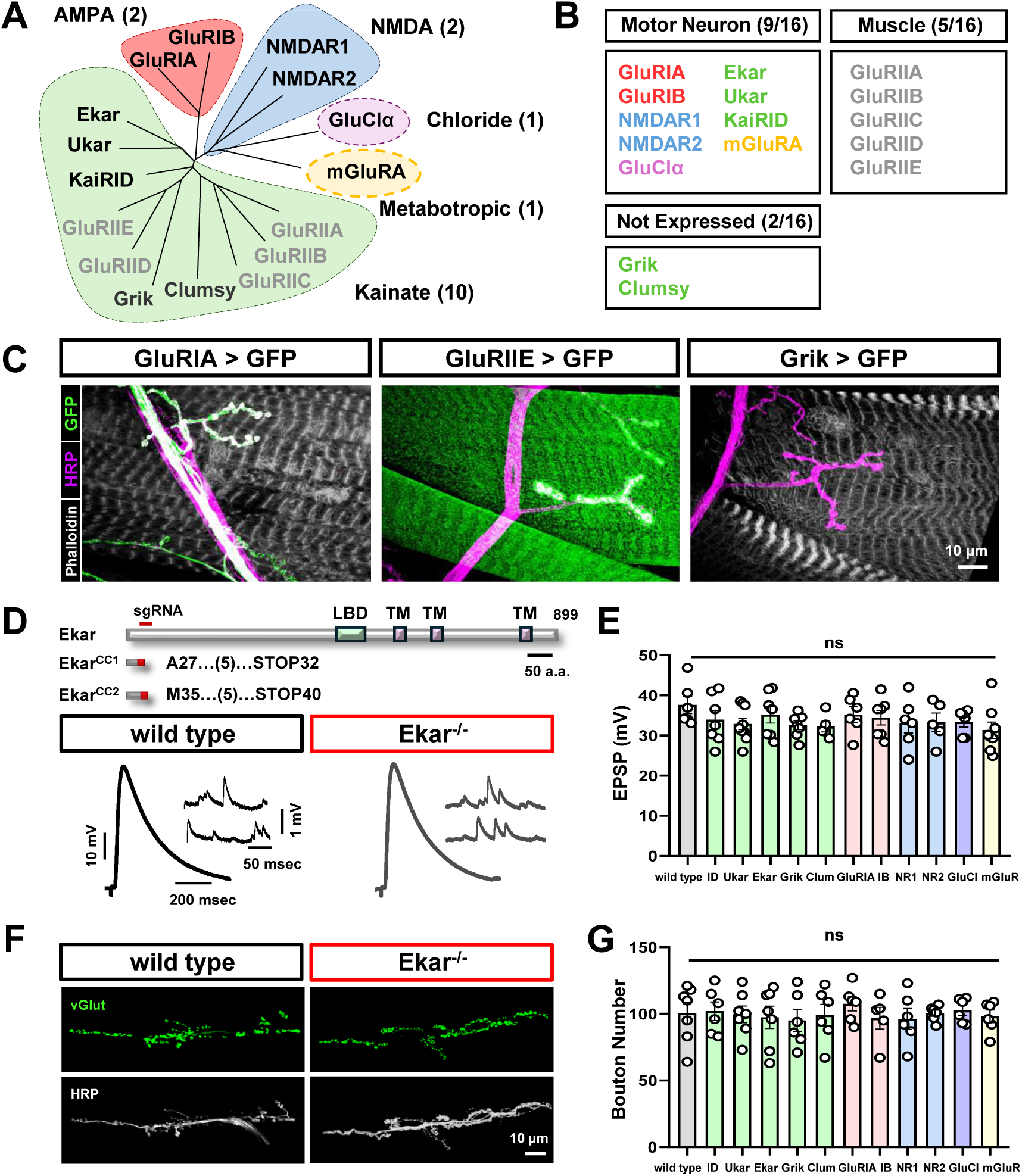
Expression mapping and CRISPR mutagenesis of all 16 GluRs encoded in the Drosophila genome. **(A)** Phylogenetic analysis, generated using the longest isoform amino acid sequence, mapping all 16 GluRs encoded in the *D. melanogaster* genome. All major GluR subtypes (AMPA, NMDA, Kainate, Chloride, and Metabotropic) are represented. **(B)** Summary of NMJ GluR expression profiles in third-instar larvae, distinguishing between motor neuron and muscle expression. **(C)** Representative NMJ images highlighting tissue-specific GluR expression using a reporter (GluR-Gal4>UAS-CD4::tdGFP) co-stained with a neuronal membrane marker (anti-HRP) and the F-actin marker phalloidin to visualize muscle. **(D)** Schematic detailing the molecular lesions in *ekar* induced by CRISPR mutagenesis (a comprehensive list of all generated GluR mutations is provided in Table S1). Accompanying representative electrophysiological traces (EPSPs and mEPSPs) of wild-type (*w^1118^*) and *ekar^CC1^* mutant NMJs are shown below. **(E)** Quantification of EPSP amplitudes across all non-muscle GluR mutants, showing no significant baseline deviations from wild type. **(F)** Representative NMJ images of wild-type and *ekar* mutants immunostained with anti-HRP and the synaptic vesicle marker vGlut to assess morphology. **(G)** Quantification of synaptic growth reveals no significant differences in bouton numbers across any of the evaluated GluR mutant alleles. Error bars indicate ±SEM. Comprehensive statistical details are available in Table S1.

We then focused on the third-instar larval NMJ. As expected, five kainate-type GluRs (GluRIIA, GluRIIB, GluRIIC, GluRIID, and GluRIIE) were exclusively expressed in the muscle (Schuster et al, 1991, Petersen et al, 1997, Marrus et al, 2004, Qin et al, 2005), which we confirmed (**Fig. 1B,C**). Critically, of the remaining five kainate receptors, three (KaiRID, Ukar, and Ekar) are expressed both in the larval brain and motor neurons (**Fig. 1B,C**), while the remaining two, Grik and Clumsy, are apparently absent from larval stages but present in adults (**Fig. S1A**). Notably, both Grik and Clumsy have been implicated in glutamatergic signaling within the adult visual system (Karuppudurai et al, 2014; Li et al, 2016), consistent with their adult-restricted expression. Furthermore, our mapping revealed that all non-kainate GluR subtypes (AMPA, NMDA, mGluR, and GluCl) are similarly expressed in the larval brain and motor neurons (**Fig. 1B,C**). We observed no evidence of GluR expression in peripheral glia. Thus, 14 of the 16 GluR subunits are expressed at the larval NMJ: five in muscle and nine in motor neurons.

Next, we utilized CRISPR/Cas9 gene editing to generate null mutations in all 16 GluRs within a common isogenic genetic background (*w^1118^*). Single-guide RNAs (sgRNAs) were designed to target the earliest shared exon among all known isoforms for each gene. Following mutagenesis, we sequenced putative mutants and isolated two independent lines harboring indel mutations that result in frameshifts and premature stop codons (**Fig. 1D** and **Table S1**). For GluClα, we used a loss of function mutant exhibiting a reduced conductance (Zak et al, 2024). Unexpectedly, we discovered that all non-muscle GluR mutants were viable, exhibiting normal adult behavior and fertility both as homozygotes and *in trans* over a defined deficiency (**Fig S1B**). Thus, beyond the essential muscle subunits GluRIIC, GluRIID, and GluRIIE which exhibit embryonic lethality (**Fig. S1B**), no other GluR subunit serves an essential function in Drosophila.

We subsequently performed a basic characterization of these non-muscle GluR mutants, assessing baseline synaptic transmission via electrophysiology and synaptic growth via confocal imaging. This comprehensive screen revealed no significant differences in miniature excitatory postsynaptic potential amplitude (mEPSP) or evoked transmission (EPSP) across any of the mutant lines compared to wild-type controls (**Fig. 1D,E** and **Table S1**). Likewise, we observed no significant alterations in bouton number (**Fig. 1F,G**). Together, these data demonstrate that while broadly expressed, none of the individual GluRs are strictly required for viability, basal neurotransmission, or synaptic growth at the larval NMJ.

### *ekar* selectively promotes chronic presynaptic homeostatic potentiation

Our primary objective was to determine which, if any, non-muscle GluRs control PHP expression. At the Drosophila NMJ, this plasticity manifests through two distinct induction mechanisms: rapid, “acute PHP” and sustained, “chronic PHP”. Acute PHP is induced by the pharmacological blockade of postsynaptic GluRs using the antagonist philanthotoxin (PhTx) (Frank et al, 2006). A 10 min incubation drives an irreversible diminution of mEPSPs, which is rapidly counteracted by a retrograde, compensatory enhancement of presynaptic glutamate release (quantal content), thereby preserving normal EPSP amplitudes (**Fig. 2A**). Conversely, chronic PHP arises from the congenital genetic loss of the muscle *GluRIIA* subunit. This yields persistently diminished mEPSP amplitudes throughout development, prompting a long-lasting, homeostatic enhancement in quantal content (**Fig. 2C**) (Petersen et al, 1997).

**Figure 2:**
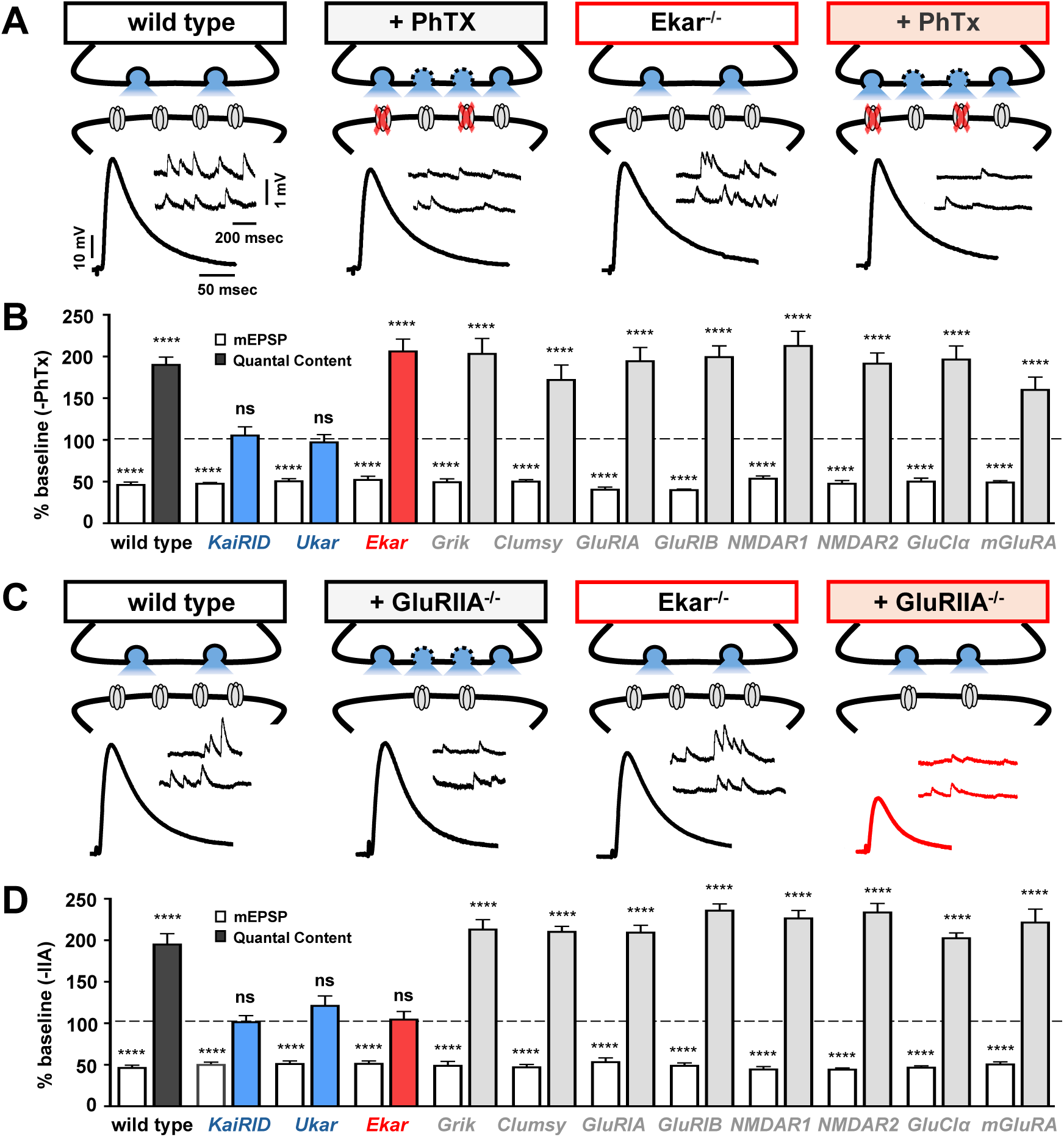
*ekar* is selectively required for chronic presynaptic homeostatic potentiation. **(A)** Schematic and representative traces of wild-type and *ekar* mutants at baseline and following acute PhTx application. **(B)** Quantification of mEPSP amplitudes and quantal content values in the indicated genotypes after PhTx treatment, normalized to untreated baseline (-PhTx) values. Consistent with previous reports, only the *kaiRID* and *ukar* mutants exhibit a robust block in acute PHP. **(C,D)** Parallel schematics, traces, and quantifications as in (A,B), evaluating chronic PHP induced by the congenital genetic loss of *GluRIIA*. Strikingly, *ekar* mutants exhibit a complete block in chronic PHP alongside the expected failures in *kaiRID* and *ukar* mutants, highlighting *ekar’s* specialized requirement for this form of plasticity. **p<0.01, ***p<0.001, ****p<0.0001; ns, not significant. See Table S1 for full statistical details.

To systematically dissect these forms of PHP, we first assessed acute PHP across the entire cohort of GluR mutants. We applied PhTx for 10 mins and compared quantal content values against untreated baselines (-PhTx). As predicted, our newly generated CRISPR alleles for *kaiRID* and *ukar* abrogated acute PHP (Kiragasi et al, 2017; Nguyen et al, 2024); despite experiencing the expected ∼50% reduction in mEPSP size, these mutants failed to mount a compensatory increase in quantal content. Crucially, extending this analysis to the remaining receptors demonstrated that every other GluR mutant – including *ekar* – exhibited robust, wild-type levels of acute PHP expression (**Fig. 2A,B**). This firmly establishes that beyond the known roles of *kaiRID* and *ukar*, no additional neuronal GluRs are deployed during the rapid induction of PHP.

Next, we interrogated the genetic requirements for chronic PHP by introducing all GluR alleles into a *GluRIIA* mutant background. This parallel approach recapitulated the expected deficits in both *kaiRID* and *ukar* mutants. However, it also unveiled a striking requirement for a previously unimplicated kainate receptor subunit, *ekar* (eye-enriched kainate receptor; Hu et al, 2015; Karuppudurai et al, 2014). Unlike its dispensability during acute plasticity, two independent *ekar* mutant alleles completely failed to express chronic PHP (**Fig. 2C,D** and Table S1). This reveals a remarkable division of labor: in contrast to the broad requirements for *kaiRID* and *ukar* in both phases, *ekar* is exclusively dedicated to chronic PHP. This selectivity strongly supports a specialized role for *ekar* in consolidating or maintaining long-term synaptic adaptations. Ultimately, this systematic genetic screen isolates Ekar as a novel executor of homeostasis and definitively limits the presynaptic receptome requirement for PHP to just three subunits: KaiRID, Ukar, and Ekar.

### *ekar* genetically interacts with *kaiRID*

The functional relationship between *ekar*, *kaiRID*, and *ukar* remains unclear. While KaiRID can form homomeric channels in heterologous systems (Li et al, 2016), and the basic phenotypes of *kaiRID* and *ukar* mutants are superficially similar in that both are required for acute and chronic PHP, *ekar* mutants expose a distinct requirement exclusively for chronic PHP. To probe these relationships *in vivo*, we performed a series of genetic interaction analyses.

A previous study demonstrated that under highly lowered extracellular Ca^2+^ conditions (0.2 mM), baseline synaptic transmission is significantly reduced in *kaiRID* mutants compared to wild-type controls (Kiragasi et al, 2017). We confirmed this deficit for both our new *kaiRID* and *ukar* alleles (**Fig. 3A,B**). Interestingly, *ekar* mutants also shared this phenotype (**Fig. 3A,B**). If *ekar* functioned through a compensatory mechanism or through a distinct genetic pathway to support basal neurotransmission, one would predict that double mutants incorporating *ekar* with *kaiRID* or *ukar* would yield a further decrement in transmission. Conversely, if *ekar* operates within the same genetic pathway, transmission should not be further reduced in the double mutants. Our epistasis experiments confirmed the latter: low Ca^2+^ recordings in double mutants (*kaiRID,ukar* and *kaiRID*;*ekar*) revealed synaptic transmission did not significantly deviate from the individual mutants alone (**Fig. 3A,B**). This confirms that all three kainate receptor subunits operate in the same genetic pathway to maintain baseline transmission under limiting Ca^2+^.

**Figure 3:**
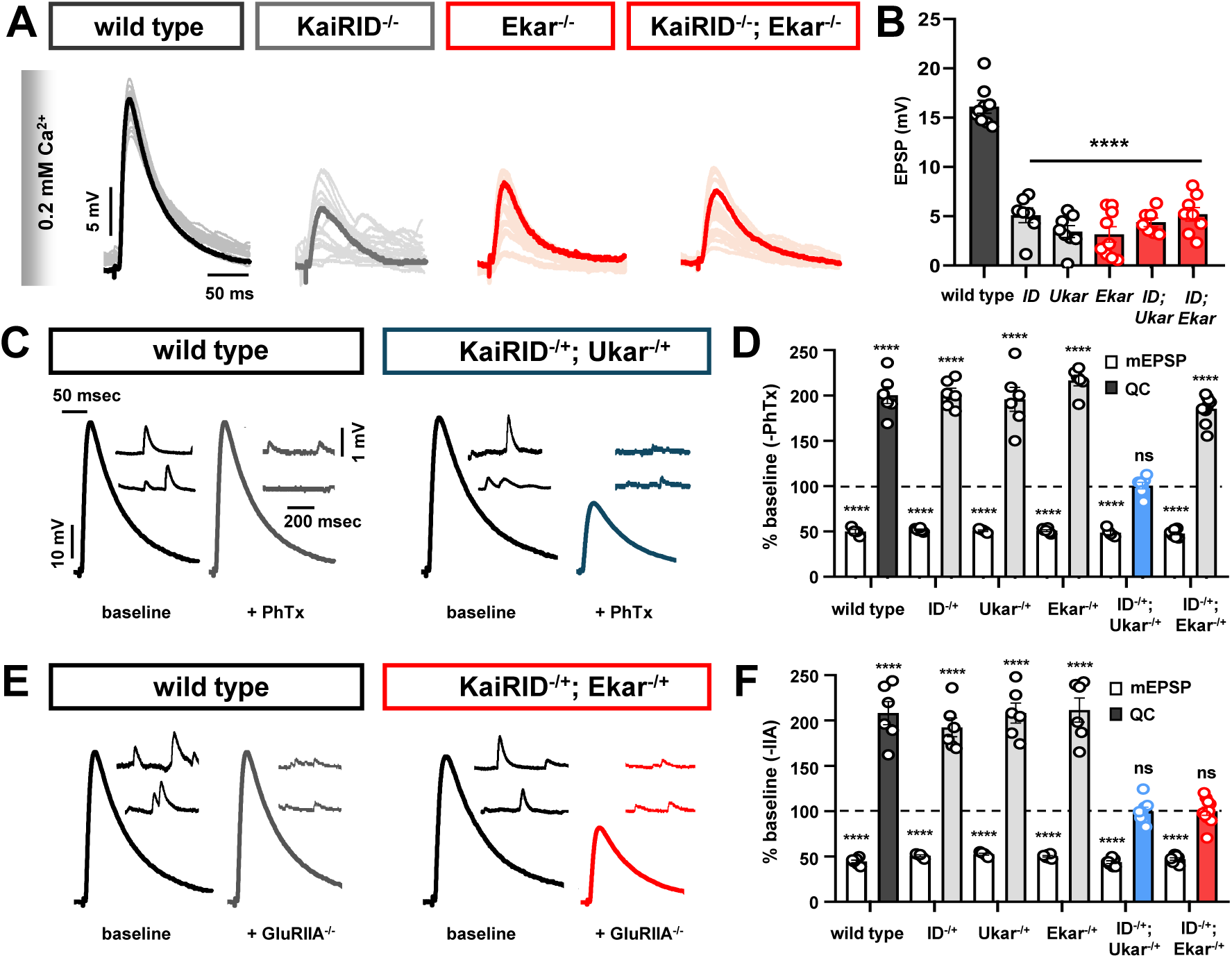
*ekar* genetically interacts with the kainate autoreceptor *kaiRID*. **(A)** Representative EPSP traces recorded under lowered external Ca^2+^ concentrations (0.2 mM). This stringent condition unmasks a deficit in baseline synaptic strength in individual GluR mutants. EPSP amplitude is not further reduced in the indicated double mutants, demonstrating that *ekar* functions in the same genetic pathway as *kaiRID* and *ukar* to promote basal neurotransmission at limiting Ca^2+^ levels. **(B)** Quantification of EPSP amplitudes across all indicated genotypes, including *kaiRID*;*ukar* and *kaiRID*;*ekar* double mutants. **(C)** Representative electrophysiological traces from wild-type and double-heterozygous (*kaiRID^−/+^*;*ukar^−/+^*) recordings at baseline and after 12 min PhTx incubation. **(D)** Quantification of mEPSP amplitude and quantal content normalized to baseline conditions (-PhTx) for the indicated genotypes. **(E)** Representative electrophysiological traces from wild-type and double-heterozygous (*kaiRID^−/+^*;*ekar^−/+^*) mutants at baseline and within a *GluRIIA* mutant background. **(F)** Quantification of mEPSP amplitudes and quantal content normalized to baseline conditions (without *GluRIIA*) for the indicated genotypes. **p<0.01, ***p<0.001, ****p<0.0001; ns, not significant. See Supplemental Table S1 for full statistical details.

Next, we executed trans-heterozygous interaction analyses, which leverage a sensitized background where the loss of a single gene copy is insufficient to disrupt a biological process. However, simultaneously removing one copy of two genes that function together in a complex or pathway can trigger a mutant phenotype, indicating a functional genetic interaction. As expected given their individual requirements for acute PHP, *kaiRID* and *ukar* double heterozygotes exhibited a complete block in acute PHP, whereas the corresponding single heterozygotes did not (**Fig. 3C,D**). Consistent with *ekar* lacking a role in acute plasticity, combining *ekar* with *kaiRID* heterozygotes did not alter acute PHP expression (**Fig. 3C,D**). In stark contrast, when assessing chronic plasticity in a *GluRIIA* mutant background, both *ukar* and *ekar* trans-heterozygotes combined with *kaiRID* yielded a severe block in chronic PHP (**Fig. 3E,F**). Together, these data provide compelling genetic evidence that all three kainate receptor subunits function within the same genetic pathway to enable chronic PHP.

### *ekar* potentiates presynaptic Ca^2+^ influx to enable chronic PHP

In our final set of experiments, we sought to clarify at which point in the homeostatic adaptation cascade *ekar* functions. In chronic PHP, retrograde signaling from *GluRIIA* mutant postsynaptic muscle triggers a robust remodeling of active zone structures, characterized by an increased abundance of the active zone scaffolds BRP (Bruchpilot), RBP (Rim Binding Protein), Unc13, and Ca_V_2/CAC channels (Weyhersmüller et al, 2011; Goel et al, 2017; Goel et al, 2019; Böhme et al, 2019). Coincident with this structural remodeling, an essential increase in presynaptic Ca^2+^ influx is observed (Müller and Davis, 2012; Chien et al, 2025; Chen et al, 2026), which ultimately drives the homeostatic increase in transmitter release. A previous study proposed that KaiRID functions as an autoreceptor during PHP, responding to glutamate release to adjust local neuronal membrane voltage (Kiragasi et al, 2017); it was hypothesized that this, in turn, drives the enhanced Ca^2+^ influx. Given this key mechanistic framework and our data establishing that *ekar* and *kaiRID* function in the same pathway, we hypothesized that *ekar* is specifically required for the homeostatic increase in Ca^2+^ influx that underpins chronic PHP.

First, we tested whether active zone remodeling persists in the absence of *ekar*. We analyzed the active zone scaffolds BRP and RBP in wild type, *GluRIIA* mutants, *ekar* mutants, and *ekar*,*GluRIIA* double mutants. In *GluRIIA* mutant controls, we observed the expected increase in fluorescence intensity compared to wild type (**Fig. 4A,B**). Crucially, in *ekar* mutants, we observed this same extent of remodeling when combined with the *GluRIIA* mutation (**Fig. 4A,B**). These data indicate that *ekar* is dispensable for the active zone remodeling cascade induced during chronic PHP.

**Figure 4:**
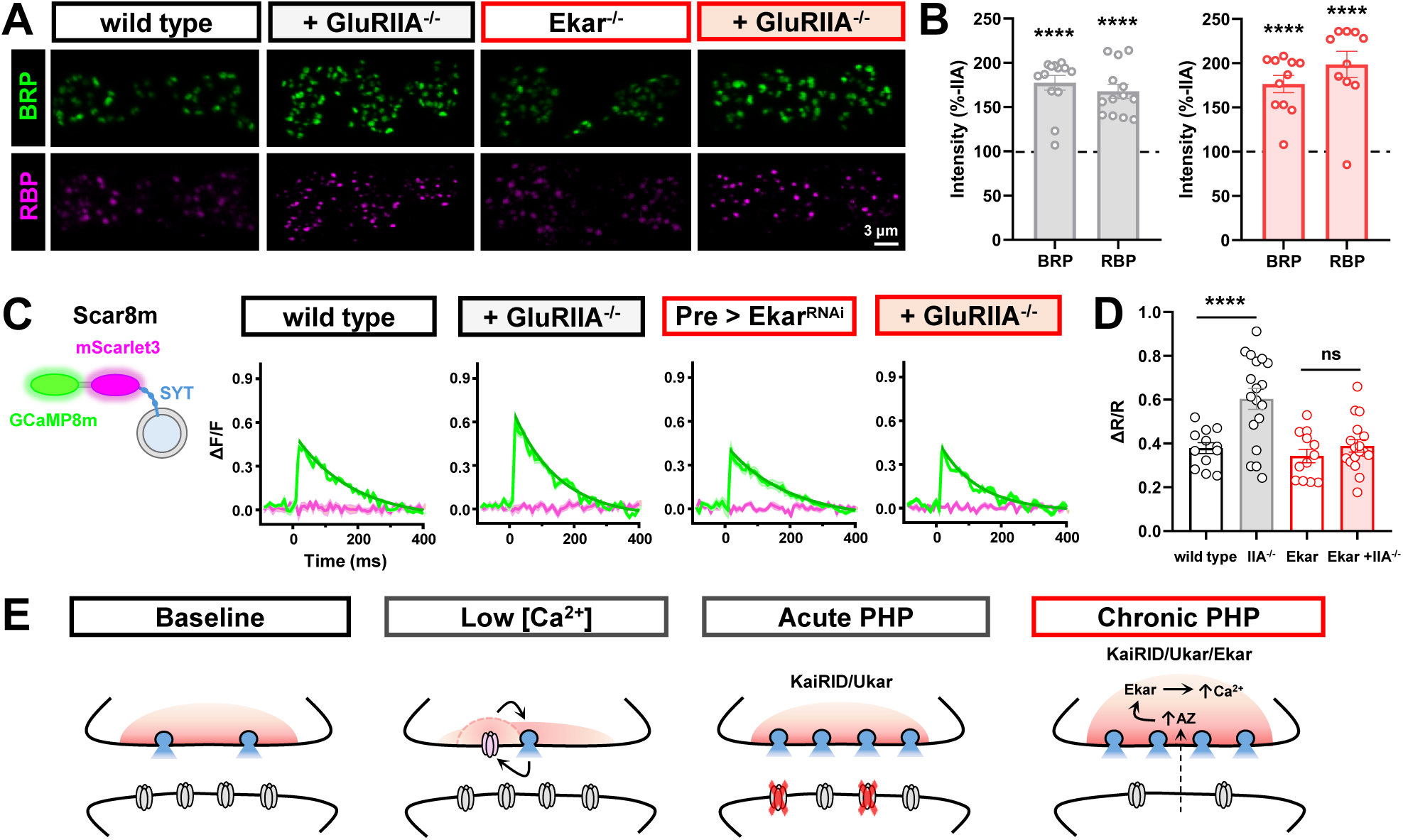
*ekar* potentiates presynaptic calcium influx to enable chronic PHP. **(A)** Representative images of MN-Ib NMJ terminals immunostained for the active zone scaffolds BRP and RBP in wild-type and *ekar* mutants at baseline and within a *GluRIIA* mutant background. Notably, both active zone components successfully remodel in *ekar*,*GluRIIA* double mutants. **(B)** Quantification of BRP and RBP mean fluorescence intensity following *GluRIIA*-induced remodeling, normalized to baseline levels of matched wild-type and *ekar* null NMJs. **(C)** Schematic of the genetically encoded presynaptic Ca^2+^ reporter utilized (“Scar8m”), which fuses the synaptic vesicle protein Synaptotagmin to GCaMP8m and mScarlet3 reporters (*OK319*>*UAS-Syt::mScarlet3::GCaMP8m*). Accompanying average ΔF/F traces illustrate presynaptic Ca^2+^ influx evoked by single action potentials. Baseline Ca^2+^ levels are comparable between wild-type and motor-neuron *ekar* knockdown NMJs (*OK319*>*ekar^RNAi^*). However, the expression of the homeostatic enhancement of Ca^2+^ influx typically observed in *GluRIIA* mutants is abolished in *ekar^RNAi^*,*GluRIIA* double-mutant NMJs. **(D)** Quantification of the ratiometric Ca²⁺ signals (ΔR/R) for each genotype. **(E)** Working model summarizing the proposed roles of presynaptic kainate receptors across four functional contexts. At baseline, presynaptic receptors are present but not actively engaged. Under low extracellular Ca^2+^ conditions, KaiRID and Ukar support baseline release probability. During acute PHP, KaiRID and Ukar drive the rapid, compensatory enhancement of quantal content. During chronic PHP, a retrograde signal from the GluRIIA-deficient muscle triggers active zone remodeling; Ekar then functions downstream of this remodeling to drive the homeostatic enhancement of presynaptic Ca^2+^ influx, which is selectively required for chronic PHP expression. See Supplemental Table S1 for full statistical details. ****p<0.0001; ns, not significant.

Next, we directly assayed presynaptic Ca^2+^ levels. To achieve this, we employed a recently engineered, ratiometric indicator, Scar8m (**Fig. 4C**) (Chen et al, 2026). We first confirmed that neuronal RNAi knockdown of *ekar* (OK319>ekar^RNAi^) sufficiently blocks chronic PHP (**Table S1**), validating this approach for Ca^2+^ imaging. We then confirmed that basal Ca^2+^ levels were indistinguishable between wild-type and ekar^RNAi^ synapses (**Fig. 4C,D**). As expected, we observed the characteristic ∼50% increase in Ca^2+^ levels in *GluRIIA* mutants (**Fig. 4C,D**). Strikingly, this homeostatic enhancement was completely abolished in *ekar* knockdown NMJs in a *GluRIIA* mutant background (**Fig. 4C,D**). Together, these findings definitively demonstrate that *ekar* functions downstream of active zone remodeling to execute the homeostatic enhancement of presynaptic Ca^2+^ influx. We propose that *ekar* likely functions as an excitatory autoreceptor to modulate presynaptic voltage and homeostatically tune Ca^2+^ entry during chronic PHP. A summary of this proposed mechanism is presented in (**Fig. 4E**).

## DISCUSSION

Through a systematic, *in vivo* CRISPR/Cas9 screen of the entire Drosophila glutamate receptome, we have generated a definitive functional and spatial atlas of the GluR gene family. This comprehensive approach yields several major conceptual advances. First, by mapping and mutating all 16 GluR subunits, we unambiguously establish the complete receptome at the neuromuscular junction, effectively ruling out previously hypothesized roles for other uncharacterized receptors in presynaptic homeostasis. Second, our screen reveals a surprising resilience in the fly genome: outside of the essential muscle GluRII subunits required for larval viability, none of the other neuronal or presynaptic GluRs appear to possess individually essential functions for organismal survival or fertility. Finally, we isolate the specific presynaptic glutamate receptor machinery required for PHP to just three kainate receptor (KAR) subunits: *kaiRID*, *ukar*, and the newly identified *ekar*. While *kaiRID* and *ukar* are broadly required for both acute and chronic PHP expression (Kiragasi et al, 2017; Nguyen et al, 2024), *ekar* is selectively essential for the long-term, chronic expression of this plasticity. Together, these findings provide a definitive molecular baseline for the Drosophila glutamate receptome and highlight a previously unrecognized division of labor among presynaptic kainate receptors in stabilizing long-term synaptic plasticity.

The precise stoichiometry and *in vivo* assembly of these three kainate receptor subunits remain an open and compelling question. Because these three KAR subunits have thus far resisted successful heterologous expression, we do not yet know whether KaiRID, Ukar, and Ekar assemble into obligate heteromeric complexes, function as distinct homomeric channels, or form a combination of subcomplexes to compartmentalize and serve specialized functions. Notably, KaiRID can form functional homomeric complexes *in vitro* (Li et al, 2016), and is antagonized by NMDA *in vitro* and *in vivo* (Li et al, 2016; Kiragasi et al, 2017; Kiragasi et al, 2020). Importantly, acute and chronic PHP target distinct inputs for homeostatic modulation: acute PHP selectively adapts release properties at phasic MN-Is, while chronic PHP targets tonic MN-Ib (Chien et al, 2025; Newman et al, 2017). It is possible, therefore, that KaiRID, Ukar, and Ekar assemble into heteromeric complexes exclusively at MN-Ib terminals to uniquely mediate chronic PHP, whereas KaiRID and Ukar may form distinct receptor complexes at MN-Is terminals to drive acute PHP. Alternatively, these subunits might function as independent subcomplexes that converge on a shared downstream effector to tune synaptic strength. Furthermore, it remains unclear how these core subunits physically associate with the essential auxiliary proteins dSol-1 and Neto-α *in vivo* (Kiragasi et al, 2020; Han et al, 2020). Given the diverse functional requirements for these subunits, one intriguing possibility is that they selectively associate with distinct auxiliary proteins to diversify their channel properties. Alternatively, they may form “tripartite” complexes encompassing core KAR subunits alongside both dSol-1 and Neto-α. This tripartite architecture is strongly supported by studies in *C. elegans*, which demonstrated that native glutamate receptors physically associate with both SOL-1 and NETO auxiliary proteins to precisely modulate channel gating and synaptic clustering (Zheng et al, 2004; Walker et al, 2006; Wang et al., 2012). Resolving the precise subunit composition and spatial distribution of these KAR complexes across distinct motor inputs will be an important next step toward understanding how input-specific homeostatic plasticity is achieved at the molecular level.

Mechanistically, how do these presynaptic KARs drive homeostatic potentiation? Previous structure-function analyses of KaiRID revealed a critical bifurcation in its ion permeability requirements. Specifically, Ca^2+^ influx through the autoreceptor is strictly required to promote baseline transmission under low-Ca^2+^ conditions (Kiragasi et al, 2017), providing a supplemental source of Ca^2+^ near release sites to boost basal release (**Fig. 4E**). Our findings suggest that Ekar and Ukar operate similarly, as their mutants share this same low-Ca^2+^ transmission deficit (**Fig. 3 A,B**). This baseline function closely mirrors presynaptic kainate autoreceptors in the mammalian brain, which facilitate transmitter release to finely tune network excitability (Schmitz et al, 2001; Pinheiro et al, 2007; Lerma and Marques 2013; Pinheiro and Mulle 2008). Intriguingly, mammalian presynaptic KARs at hippocampal mossy fiber synapses also operate through dual signaling modes: a Ca^2+^-dependent, ionotropic mechanism that facilitates baseline release, and a distinct, metabotropic-like pathway that modulates release independently of ion flux (Pinheiro and Mulle 2008; Lerma and Marques 2013). This duality closely parallels the bifurcated ion permeability requirements of KaiRID at the Drosophila NMJ, where Ca^2+^ conductance supports baseline neurotransmission but sodium-driven depolarization drives PHP (Kiragasi et al, 2017). However, Ca^2+^ permeability through KaiRID is entirely dispensable for the expression of PHP. Instead, sodium-driven membrane depolarization through these receptors likely modulates the presynaptic action potential waveform to drive PHP (Kiragasi et al, 2017). This mechanism is highly analogous to the presynaptic DEG/ENaC channel PPK1/11/16, which regulates PHP by modulating presynaptic voltage to drive a subsequent, downstream enhancement in Ca^2+^ influx through Ca_V_2/CAC channels (Younger et al, 2013; Orr et al, 2017). Collectively, our data support a model in which Ekar-containing KAR complexes function as excitatory autoreceptors that modulate presynaptic voltage, linking retrograde PHP signaling to the enhanced Ca^2+^ influx specifically required for chronic PHP expression (**Fig. 4E**).

The identification of this highly specialized presynaptic KAR complex at the Drosophila NMJ reveals striking conceptual parallels with the homeostatic regulation of mammalian central synapses. In the mammalian brain, elegant work has demonstrated that the auxiliary subunit Neto2 and its associated KARs are critically required for postsynaptic homeostatic receptor scaling (Yan et al, 2013). Our findings extend and invert this synaptic paradigm, demonstrating that Drosophila KARs function on the presynaptic side to homeostatically regulate synaptic strength. Moreover, like their mammalian counterparts, these fly KARs require auxiliary subunits – dSol-1 and Neto-α – for their function (Kiragasi et al, 2020, Han et al, 2020), suggesting deep evolutionary conservation of the KAR-auxiliary protein partnership in homeostatic signaling. Ultimately, this establishes a deeply conserved framework where KARs and their associated auxiliary subunits function as compartmentalized homeostatic regulators, poised on both sides of the synapse to maintain stable neural circuit function. The complete genetic toolkit and definitive receptome atlas established here now provide the foundation to determine how distinct KAR subunit combinations are assembled, trafficked to specific release sites, and differentially regulated to achieve the input-specific homeostatic plasticity that maintains stable circuit function throughout the lifetime of the organism.

## MATERIALS AND METHODS

### Fly stocks

All Drosophila stocks were maintained at 25°C on standard molasses food. The *w^1118^* strain served as the wild-type control and the isogenic background for all experimental genotypes. All experiments used third-instar larvae of both sexes. A comprehensive list of fly stocks and their sources is available in Table S2.

### Molecular Biology

GluR mutants were generated using a CRISPR/Cas9 gene-editing strategy as previously described (Stark et al, 2025). Single-guide RNA (sgRNA) sequences (listed in Table S1) were designed to target the earliest shared exons across all known isoforms of each respective gene. Two to three sgRNAs per gene were cloned into the pU63 vector (Addgene #112811), incorporating intervening tRNA (F + E) sequences to facilitate multiplexed gRNA expression. Constructs were injected by BestGene (Chino Hills, CA) for targeted insertion into the attP2 or attP40 docking sites. Transgenic gRNA lines were subsequently crossed to *Bam-Cas9* (BDSC 97543) or *Nos-Cas9* (BDSC 78782) to induce germline CRISPR mutagenesis. PCR screening of 20 independent lines identified two distinct indel alleles per target that successfully shifted the open reading frame and introduced premature stop codons, confirming them as null mutants. To eliminate potential off-target effects, the gRNA and Cas9 transgenes were restricted to the germline for a single generation prior to outcrossing to the isogenic *w^1118^* background.

A T2A-Gal4 knock-in allele of the Drosophila glutamate receptor gene Ekar was generated using PhiC31-mediated recombination-mediated cassette exchange (RMCE) (WellGenetics Inc., Taipei City, Taiwan). The donor construct, pBS-KS-attB2-SA(1)-T2A-Gal4-Hsp70 (Addgene #62897), was co-injected with PhiC31 integrase into embryos of the MiMIC line *Ekar^MI09564^* (BDSC 60827). Site-specific cassette exchange at the MiMIC insertion (MI09564) replaced the original cassette with a splice acceptor–T2A–Gal4 sequence, enabling Gal4 expression under endogenous *ekar* regulatory control. F1 progeny were screened for successful RMCE events by the loss of the yellow marker (y−), followed by confirmation via genomic PCR and sequencing. The resulting allele, *ekar^MI09564-Trojan-T2A-Gal4.1^*, provides a robust genetic tool for driving Gal4 expression in *ekar*-expressing cells.

### Electrophysiology

Sharp-electrode current clamp recordings were performed on abdominal muscle 6 or 7 in segments A3 or A4 of third-instar larvae following established procedures (Qiu et al, 2025). Preparations were dissected and recorded in modified hemolymph-like saline (HL-3) containing (in mM): 70 NaCl, 5 KCl, 10 MgCl_2_, 10 NaHCO_3_, 115 Sucrose, 5 Trehelose, 5 HEPES, and 0.5 CaCl_2_ (pH 7.2), unless otherwise stated. Only cells with an initial resting potential below −60 mV and an input resistance greater than 5 MΩ were included for analysis. Spontaneous miniature excitatory postsynaptic potentials (mEPSPs) were recorded for 60 secs without stimulation and low-pass filtered at 1 kHz. Data were acquired using an Axoclamp 900A amplifier, a Digidata 1440A system, and pClamp 10.5 software (Molecular Devices) on an Olympus BX61 WI microscope with a 40x/0.80 NA water-dipping objective. mEPSP parameters were analyzed using MiniAnalysis (Synaptosoft) and Excel (Microsoft), with the average amplitude per NMJ derived from approximately 100 events. To induce acute postsynaptic receptor blockade, dissected larvae were incubated in 20 µM philanthotoxin-433 (PhTx; Sigma) in HL-3 for 12 mins prior to recording.

### Calcium Imaging

Live presynaptic Ca^2+^ imaging was performed on third-instar larval NMJs expressing the Scar8m sensor (*OK319*>*UAS-Syt::mScarlet3::GCaMP8m*) (Chen et al, 2026). Preparations were imaged using an Olympus BX61WI upright microscope equipped with a 40×/0.80 NA water-dipping objective and a Hamamatsu camera controlled via NIS-Elements software. GCaMP and mScarlet signals were simultaneously captured using a single dual-emission filter under blue and green LED illumination (25% power each). Fast time-lapse acquisitions were recorded at 100 frames per second. Motor neurons were electrically stimulated at 1 Hz (1-msec pulse duration), capturing four evoked Ca^2+^ events per recording. Regions of interest (ROIs) were manually defined around three MN-Ib terminal boutons on muscle 6/7 of abdominal segment A3. Fluorescence intensity changes (ΔF/F) across frames were quantified using NIS-Elements software. The ratiometric intensity changes (ΔR/R) were quantified using Cafire program (Chen et al, 2026). A minimum of five animals per genotype were analyzed.

### Immunocytochemistry

Third-instar larvae were dissected in ice-cold Ca^2+^-free HL-3 (Perry et al, 2022). Preparations were fixed in Bouin’s solution, ice-cold methanol, or 4% paraformaldehyde, followed by a 30 min wash in PBST (0.1% Triton X-100 in PBS) and blocking in 5% normal donkey serum. Samples were incubated with primary antibodies overnight at 4°C. Following three PBST washes, preparations were incubated with secondary antibodies for 2 hours at room temperature in the dark, washed three additional times in PBST, and equilibrated in 70% glycerol. Samples were mounted in VectaShield (Vector Laboratories: H-1000-10) for imaging.

Adult Drosophila brains were dissected in ice-cold phosphate-buffered saline (PBS) and immediately fixed in 4% paraformaldehyde (PFA) for 20 minutes at room temperature. Following fixation, tissues were washed six times in 0.3% Triton X-100 in PBS (PBST) and subsequently blocked in 5% normal donkey serum (NDS) for 1 hour. Brains were then incubated with primary antibodies diluted in blocking buffer overnight at 4°C. After extensive washing in PBST (three cycles of six washes each, with 15-minute room temperature incubations), tissues were incubated with secondary antibodies for 2 hours at room temperature, shielded from light. Following final PBST washes, brains were mounted in Vectashield for confocal imaging. Antibody sources, dilutions, and references are detailed in Table S2.

### Confocal imaging and analysis

NMJs were imaged using a Nikon A1R Confocal microscope equipped with a 100x APO 1.4 NA oil immersion objective and NIS Elements software (Han et al, 2023). Images were acquired across four separate laser lines (405 nm, 488 nm, 561 nm, and 647 nm). To accurately quantify BRP and RBP fluorescence intensity, all compared genotypes were imaged on the same day using identical laser power and detector gain settings. Z-stacks were acquired with a 0.150 μm step size and an x/y pixel size of 40 nm. Raw images were deconvolved using Huygens Object Analyzer (SVI) to identify individual puncta and quantify mean fluorescence intensities. Intensity analyses were restricted to MN-Ib terminals of muscle 6/7 abdominal segment A3. All metrics were derived from a minimum of five animals per genotype.

### Statistical analysis

Statistical analyses were performed using GraphPad Prism (version 7.0), MiniAnalysis, and Microsoft Excel. Data were first evaluated for normal distribution using the D’Agostino & Pearson omnibus test, followed by a one-way ANOVA with a post-hoc Tukey’s multiple comparisons test to assess significance between groups. All figures present data as mean ±SEM. Statistical significance is denoted as: p<0.05 (*), p<0.01 (**), p<0.001 (***), and p<0.0001 (****), with ‘ns’ indicating non-significant differences. Comprehensive statistical details and sample sizes (n) are provided in Table S1.

## ACKNOWLEDGEMENTS

We acknowledge the Developmental Studies Hybridoma Bank (Iowa, USA) for antibodies used in this study, and the Bloomington Drosophila Stock Center (NIH P40OD018537) for fly stocks. This work was supported by a grant from the National Institutes of Health (NS091546) and NSF grant IOS-2417451 to DD and NSF-BSF Grant 2024612 to G.S.O. J.M. was supported by an NIH supplement to the same grant.

## AUTHOR CONTRIBUTIONS

JM, CC, and DD designed the research, JM, CC, WD, AC, and SS performed experiments, and JM and CC analyzed the data; JM, CC, and NT generated CRISPR alleles; HZ and GSO contributed reagents. The manuscript was written by JM and DD with feedback from the other authors.

---

**Supplemental Figure S1: GluR expression in the larval and adult brains. (A)** Tabular summary of GluR expression detected in the larval and adult Drosophila central nervous systems. **(B)** Summary of viability and lethal stages for all generated GluR null mutants (V=viable, E=embryonic lethal). **(C)** Representative confocal images of larval brain GluR expression visualized via a GFP reporter (GluR-Gal4>UAS-CD4::tdGFP) and co-stained with a neuronal membrane marker (HRP). **(D)** Representative images of adult brain GluR expression highlighting distinct regional localization patterns.

