## Supplemental Figure 1, Supplemental Table 1, Supplemental Table 2 for "A comprehensive CRISPR screen of the Drosophila glutamate receptome reveals Ekar as a selective regulator of presynaptic homeostatic plasticity"

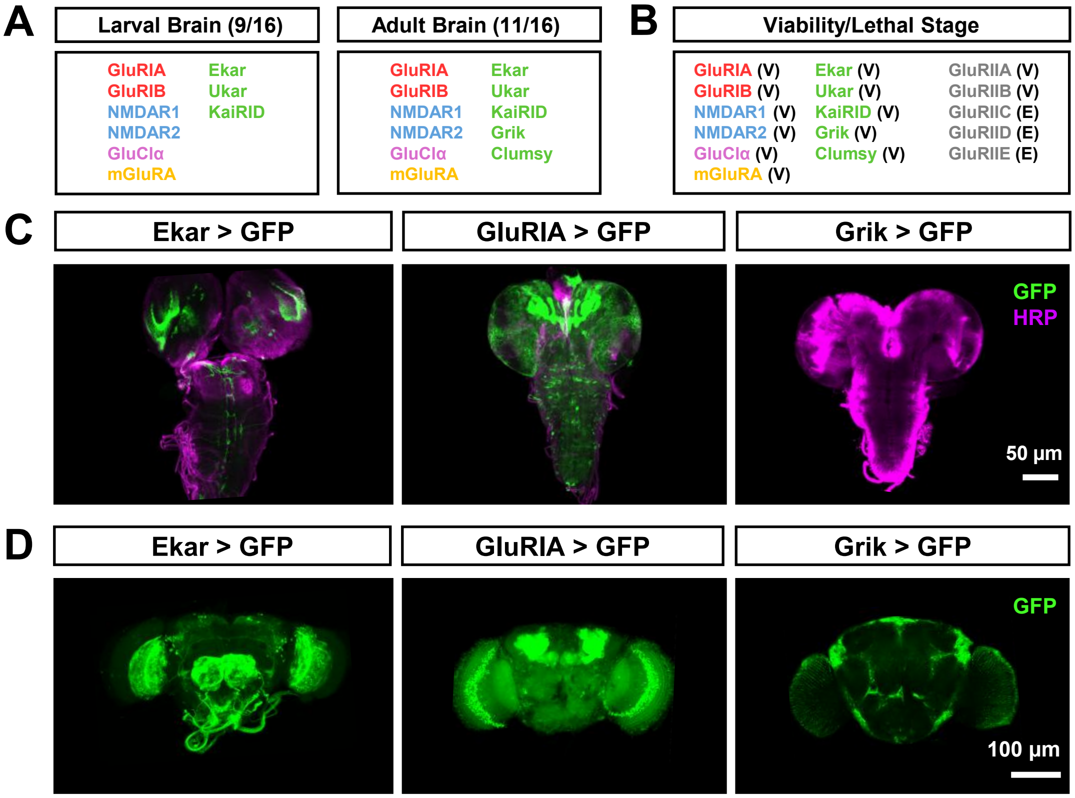


**Figure S1: GluR expression in the larval and adult brains.**

**Supplementary Table 1: Absolute values for normalized data and additional statistical details.** The figure and panel, genotype, and conditions are noted. Average values (with standard error of the mean noted in parentheses), data samples (n), and statistical significance tests are shown for all data.

| Figure | GluR | gRNA 1 | gRNA 2 | gRNA 3 | Landing site | Allele 1 Molecular Lesion | Allele 2 Molecular Lesion |
|---|---|---|---|---|---|---|---|
| 1D | KaiRID | TGATGCGGTCCACC GCCTGG | CCGGCATGGAAGCT GTCGAA | - | attp40 | R49…(37)…STOP86 | - |
| 1D | Ukar | AGTGCGAAACGAGT CATCTC | GGTCAATTCGACCT TCAATA | - | attp40 | L33...(3)...STOP36 | V34…(12)…STOP46 |
| 1D | Ekar | GTGTGCAAACTTAT ACGAGT | TGGGCAGACCGGC CTATGTT | - | attp40 | A27…(5)…STOP32 | M35A…(5)…STOP40 |
| 1D | Grik | GATCTTCTCGAGAT CCGGAT | TCACGCACAGCAAA ATCTGG | - | attp40 | K84…(19)…STOP103 | V72…(67)…STOP139 |
| 1D | Clumsy | GATAACCTCTTGTT GTCCCA | GGGGCTGGCGGAG ATTGTGC | GGTCGAGGCCAAC ATAACTA | attp2 | S70…(3)…STOP73 | L68…(46)…STOP114 |
| 1D | GluRIA | TGCAATCTTCGAAC AGGGAA | CTCTCGACGCTTCG AATTGC | CGCGGAGTCTACTC AATGCT | attp40 | G49…(3)…STOP52 | S99…(21)…STOP120 |
| 1D | GluRIB | GGCGGTGGAGGTA GCGGGGT | TCGTGACGCCCTGG TTCCCC | CGACTTTGCCCTCA GCATGC | attp40 | P151…(13)…STOP164 | I152…(16)…STOP168 |
| 1D | NMDAR2 | AAACTGAAACGCGG CACGGA | ATCTGGTGTCCAGG ATCAAC | GTCGCTGACGCCCA GCCCGA | attp2 | P70…(8)… STOP78 | G11…(31)…STOP42 |
| 1D | mGluRA | TAATTCCAGGAAGT ATGTTG | CTAAACAATTTAGA CACATC | - | attp40 | V42…(14)…STOP56 | C40…(40)…STOP80 |

| Figure | Label | Genotype | $[Ca^{2+}]_E$ (mM) | mEPSP amplitude (mV) | mEPSP frequency (Hz) | EPSP amplitude (mV) | QC | Rinput (MΩ) | Resting potential (mV) | n | P Value (significance: mEPSP, mEPSP freq., EPSP) |
|---|---|---|---|---|---|---|---|---|---|---|---|
| 1E | wild type | $w^{1118}$ | 0.4 | 1.10 (±0.07) | 3.36 (±1.79) | 36.62 (±2.16) | 32.29 (±1.96) | 9.10 (±0.21) | -68.78 (±1.81) | 6 | - |
| 1E | ID | $w;+;kaiRID^{JM1}$ | 0.4 | 0.99 (±0.03) | 2.52 (±1.88) | 35.20 (±2.11) | 35.56 (±2.13) | 8.24 (±0.35) | -69.54 (±1.54) | 7 | 0.12 (ns), 0.44 (ns), 0.65 (ns) |
| 1E | Ukar | $w;+;+;ukar^{CC1}$ | 0.4 | 1.06 (±0.06) | 3.26 (±0.56) | 32.89 (±1.44) | 31.03 (±1.36) | 10.98 (±0.32) | -63.22 (±2.65) | 9 | 0.28 (ns), 0.36 (ns), 0.33 (ns) |
| 1E | Ekar | $w;+;+;ekar^{CC1}$ | 0.4 | 1.04 (±0.07) | 3.21 (±1.52) | 33.92 (±2.27) | 32.62 (±2.19) | 11.21 (±0.15) | -65.57 (±2.97) | 7 | 0.79 (ns), 0.27 (ns), 0.53 (ns) |
| 1E | Grik | $w;+;grik^{JM1}$ | 0.4 | 1.09 (±0.06) | 3.67 (±0.79) | 32.22 (±1.31) | 29.56 (±0.65) | 9.36 (±0.41) | -65.93 (±1.77) | 7 | 0.47 (ns), 0.76 (ns), 0.20 (ns) |
| 1E | Clumsy | $w;clumsy^{NT1}$ | 0.4 | 1.13 (±0.06) | 2.85 (±1.12) | 35.23 (±1.90) | 31.18 (±1.68) | 12.81 (±0.21) | -71.98 (±0.98) | 6 | 0.99 (ns), 0.42 (ns), 0.24 (ns) |
| 1E | IA | $w;+;gluRIA^{NT1}$ | 0.4 | 1.05 (±0.09) | 2.89 (±1.60) | 34.44 (±1.81) | 32.80 (±1.72) | 8.85 (±0.34) | -71.41 (±1.36) | 6 | 0.13 (ns), 0.28 (ns), 0.52 (ns) |
| 1E | IB | $w;+;gluRIB^{NT1}$ | 0.4 | 1.06 (±0.05) | 2.96 (±1.74) | 33.16 (±2.51) | 31.28 (±2.37) | 9.73 (±0.36) | -67.08 (±1.93) | 7 | 0.44 (ns), 0.35 (ns), 0.61 (ns) |
| 1E | NR1 | $w;+;NMDAR1^{XC1}$ | 0.4 | 1.20 (±0.06) | 3.48 (±1.28) | 33.25 (±2.33) | 27.71 (±1.94) | 9.82 (±0.22) | -65.64 (±1.22) | 6 | 0.54 (ns), 0.74 (ns), 0.29 (ns) |
| 1E | NR2 | $w,NMDAR2^{NT1};+;+$ | 0.4 | 1.10 (±0.08) | 4.05 (±0.64) | 33.40 (±1.29) | 30.36 (±1.17) | 10.11 (±0.27) | -63.97 (±1.90) | 6 | 0.31 (ns), 0.71 (ns), 0.25 (ns) |
| 1E | GluCl | $w;+;GluCl\alpha^{1}$ | 0.4 | 0.97 (±0.05) | 2.45 (±1.33) | 31.35 (±2.02) | 32.32 (±2.09) | 11.01 (±0.41) | -73.05 (±1.64) | 6 | 0.53 (ns), 0.82 (ns), 0.47 (ns) |
| 1E | mGluR | $w;+;+;mGluRA^{NT1}$ | 0.4 | 1.08 (±0.08) | 3.32 (±1.67) | 32.54 (±1.10) | 30.13 (±1.02) | 10.76 (±0.24) | -70.76 (±2.01) | 7 | 0.31 (ns), 0.23 (ns), 0.39 (ns) |

| Figure | Label | Genotype | Muscle 6/7 Bouton #/NMJ | n | P Value (significance: Bouton #) |
|---|---|---|---|---|---|
| 1G | wild type | $w^{1118}$ | 100.6 (±7.93) | 7 | - |
| 1G | KaiRID | $w;+;kaiRID^{JM1}$ | 102.0 (±6.87) | 6 | 0.89 (ns) |
| 1G | Ukar | $w;+;+;ukar^{CC1}$ | 101.5 (±6.25) | 7 | 0.96 (ns) |
| 1G | Ekar | $w;+;+;ekar^{CC1}$ | 96.67 (±8.38) | 7 | 0.79 (ns) |
| 1G | Grik | $w;+;grik^{JM1}$ | 95.0 (±8.28) | 6 | 0.84 (ns) |
| 1G | Clumsy | $w;clumsy^{NT1}$ | 99.0 (±8.21) | 6 | 0.74 (ns) |
| 1G | GluRIA | $w;+;gluRIA^{NT1}$ | 107.5 (±5.34) | 6 | 0.41 (ns) |
| 1G | GluRIB | $w;+;gluRIB^{NT1}$ | 96.8 (±8.10) | 6 | 0.28 (ns) |
| 1G | NMDAR1 | $w;+;NMDAR1^{XC1}$ | 96.3 (±7.77) | 6 | 0.97 (ns) |
| 1G | NMDAR2 | $w,NMDAR2^{NT1};+ ;+$ | 100.5 (±2.33) | 6 | 0.62 (ns) |

| | | | | | |
|---|---|---|---|---|---|
| 1G | GluClα | *w;+;GluClα*$^{1}$ | 102.7 (3.73) | 6 | 0.63 (ns) |
| 1G | mGluRA | *w;+;+;mGluRA*$^{NT1}$ | 98.2 (4.71) | 6 | 0.47 (ns) |

| Figure | Label | Genotype (+PhTx) | [Ca$^{2+}$]$_E$ (mM) | mEPSP amplitude (mV) | EPSP amplitude (mV) | QC | Rinput (MΩ) | Resting potential (mV) | n | P Value (significance): mEPSP, QC |
|---|---|---|---|---|---|---|---|---|---|---|
| 2A | wild type | *w*$^{1118}$ | 0.4 | 0.48 (±0.06) | 33.56 (±2.01) | 69.92 (±3.25) | 5.27 (±0.54) | -71.76 (±2.79) | 10 | <0.0001 (****), <0.0001 (****) |
| 2A | KaiRID | *w;+;kaiRID*$^{JM1}$ | 0.4 | 0.54 (±0.03) | 20.24 (±1.61) | 37.48 (±3.24) | 8.49 (±0.47) | -66.47 (±1.82) | 8 | <0.0001 (****), 0.3695 (ns) |
| 2A | Ukar | *w;+;+;ukar*$^{CC1}$ | 0.4 | 0.51 (±0.03) | 16.45 (±1.31) | 32.25 (±3.32) | 6.18 (±1.04) | -69.34 (±1.51) | 9 | <0.0001 (****), 0.9822 (ns) |
| 2A | Ekar | *w;+;+;ekar*$^{CC1}$ | 0.4 | 0.43 (±0.04) | 30.19 (±3.34) | 70.21 (±3.72) | 10.24 (±1.24) | -64.06 (±1.77) | 9 | <0.0001 (****), <0.0001 (****) |
| 2A | Grik | *w;+;grik*$^{JM1}$ | 0.4 | 0.46 (±0.06) | 30.27 (±1.25) | 65.80 (±2.71) | 10.35 (±0.48) | -69.85 (±1.91) | 7 | <0.0001 (****), <0.0001 (****) |
| 2A | Clumsy | *w;clumsy*$^{NT1}$ | 0.4 | 0.53 (±0.06) | 31.28 (±2.19) | 60.91 (±3.85) | 6.46 (±0.57) | -64.98 (±2.67) | 7 | <0.0001 (****), <0.0001 (****) |
| 2A | GluRIA | *w;+;gluRIA*$^{NT1}$ | 0.4 | 0.46 (±0.05) | 30.96 (±2.24) | 67.30 (±3.22) | 7.55 (±0.53) | -63.71 (±3.15) | 7 | <0.0001 (****), <0.0001 (****) |
| 2A | GluRIB | *w;+;gluRIB*$^{NT1}$ | 0.4 | 0.48 (±0.06) | 31.89 (±1.73) | 66.43 (±2.32) | 4.35 (±0.89) | -69.46 (±2.84) | 7 | <0.0001 (****), <0.0001 (****) |
| 2A | NMDAR1 | *w;+;NMDAR1*$^{XC1}$ | 0.4 | 0.43 (±0.08) | 30.54 (±1.17) | 71.02 (±3.68) | 6.54 (±0.25) | -65.87 (±2.33) | 7 | <0.0001 (****), <0.0001 (****) |
| 2A | NMDAR2 | *w,NMDAR2*$^{NT1}$*;+;+* | 0.4 | 0.46 (±0.04) | 29.57 (±0.98) | 64.28 (±2.37) | 8.24 (±0.62) | -64.71 (±3.81) | 8 | <0.0001 (****), <0.0001 (****) |
| 2A | GluClα | *w;+;GluClα*$^{1}$ | 0.4 | 0.52 (±0.03) | 32.18 (±1.30) | 61.88 (±2.17) | 10.34 (±0.59) | -65.26 (±2.15) | 7 | <0.0001 (****), <0.0001 (****) |
| 2A | mGluRA | *w;+;+;mGluRA*$^{NT1}$ | 0.4 | 0.54 (±0.06) | 28.29 (±2.68) | 52.38 (±3.26) | 9.48 (±1.32) | -73.42 (±1.70) | 7 | <0.0001 (****), <0.0001 (****) |

| Figure | Label | Genotype (+GluRIIA$^{-/-}$) | [Ca$^{2+}$]$_E$ (mM) | mEPSP amplitude (mV) | EPSP amplitude (mV) | QC | Rinput (MΩ) | Resting potential (mV) | n | P Value (significance): mEPSP, QC |
|---|---|---|---|---|---|---|---|---|---|---|
| 2B | wild type | *w;GluRIIA*$^{PV3}$ | 0.4 | 0.52 (±0.06) | 32.91 (±2.48) | 63.28 (±3.83) | 8.61 (±0.23) | -65.14 (±2.19) | 11 | <0.0001 (****), <0.0001 (****) |
| 2B | KaiRID | *w;GluRIIA*$^{PV3}$*; kaiRID*$^{JM1}$ | 0.4 | 0.54 (±0.04) | 19.65 (±2.24) | 36.38 (±2.39) | 9.16 (±0.55) | -64.64 (±1.49) | 8 | <0.0001 (****), 0.2803 (ns) |
| 2B | Ukar | *w;GluRIIA*$^{PV3}$*;+; ukar*$^{CC1}$ | 0.4 | 0.51 (±0.03) | 19.32 (±2.98) | 37.88 (±3.37) | 9.45 (±0.64) | -67.15 (±2.37) | 9 | <0.0001 (****), 0.2469 (ns) |
| 2B | Ekar | *w;GluRIIA*$^{PV3}$*;+; ekar*$^{CC1}$ | 0.4 | 0.52 (±0.07) | 17.89 (±2.56) | 34.40 (±2.84) | 10.34 (±0.28) | -68.51 (±1.89) | 13 | <0.0001 (****), 0.6702 (ns) |
| 2B | Grik | *w;GluRIIA*$^{PV3}$*; grik*$^{JM1}$ | 0.4 | 0.48 (±0.04) | 30.35 (±2.67) | 62.23 (±3.21) | 5.32 (±1.03) | -68.46 (±2.36) | 7 | <0.0001 (****), <0.0001 (****) |
| 2B | Clumsy | *w;GluRIIA*$^{PV3}$*, clumsy*$^{NT1}$ | 0.4 | 0.51 (±0.03) | 33.59 (±1.49) | 65.86 (±1.69) | 7.18 (±0.91) | -65.81 (±2.57) | 7 | <0.0001 (****), <0.0001 (****) |
| 2B | GluRIA | *w;GluRIIA*$^{PV3}$*; gluRIA*$^{NT1}$ | 0.4 | 0.51 (±0.05) | 34.61 (±2.18) | 67.86 (±2.47) | 6.95 (±0.67) | -73.61 (±3.62) | 7 | <0.0001 (****), <0.0001 (****) |
| 2B | GluRIB | *w;GluRIIA*$^{PV3}$*; gluRIB*$^{NT1}$ | 0.4 | 0.46 (±0.07) | 35.79 (±1.72) | 77.80 (±2.16) | 8.17 (±0.37) | -72.73 (±1.82) | 7 | <0.0001 (****), <0.0001 (****) |
| 2B | NMDAR1 | *w;GluRIIA*$^{PV3}$*; NMDAR1*$^{XC1}$ | 0.4 | 0.54 (±0.05) | 34.02 (±2.21) | 63.00 (±2.36) | 9.67 (±0.88) | -66.27 (±2.71) | 8 | <0.0001 (****), <0.0001 (****) |
| 2B | NMDAR2 | *w,NMDAR2*$^{NT1}$*; GluRIIA*$^{PV3}$*;+* | 0.4 | 0.45 (±0.05) | 32.03 (±2.34) | 71.17 (±3.00) | 8.45 (±0.29) | -68.53 (±1.27) | 7 | <0.0001 (****), <0.0001 (****) |
| 2B | GluClα | *w;GluRIIA*$^{PV3}$*; GluClα*$^{1}$ | 0.4 | 0.53 (±0.05) | 34.84 (±1.63) | 65.73 (±1.78) | 7.62 (±0.71) | -69.18 (±2.36) | 7 | <0.0001 (****), <0.0001 (****) |
| 2B | mGluRA | *w;GluRIIA*$^{PV3}$*;+; mGluRA*$^{NT1}$ | 0.4 | 0.46 (±0.03) | 30.83 (±2.64) | 67.02 (±4.57) | 7.68 (±1.06) | -71.24 (±3.12) | 8 | <0.0001 (****), <0.0001 (****) |

| Figure | Label | Genotype | [Ca$^{2+}$]$_E$ (mM) | EPSP amplitude (mV) | Rinput (MΩ) | Resting potential (mV) | n | P Value (significance): mEPSP, QC |
|---|---|---|---|---|---|---|---|---|
| 3 A,B | wild type | *w*$^{1118}$ | 0.2 | 16.04 (±1.08) | 8.52 (±0.33) | -66.86 (±1.63) | 10 | - |
| 3 A,B | ID | *w;+;kaiRID*$^{JM1}$ | 0.2 | 5.10 (±0.75) | 10.31 (±0.99) | -68.24 (±1.68) | 7 | <0.0001 (****) |
| 3 A,B | Ukar | *w;+;+;ukar*$^{CC3}$ | 0.2 | 3.45 (±0.60) | 9.81 (±1.17) | -68.98 (±1.24) | 8 | <0.0001 (****) |
| 3 A,B | Ekar | *w;+;+;ekar*$^{CC1}$ | 0.2 | 3.16 (±0.78) | 12.45 (±0.93) | -67.81 (±2.45) | 9 | <0.0001 (****) |
| 3 A,B | ID;Ukar | *w;+;kaiRID*$^{JM1}$*;ukar*$^{CC3}$ | 0.2 | 4.39 (±0.44) | 9.62 (±0.85) | -64.21 (±1.44) | 7 | <0.0001 (****) |
| 3 A,B | ID;Ekar | *w;+;kaiRID*$^{JM1}$*;ekar*$^{CC1}$ | 0.2 | 5.20 (±0.69) | 8.46 (±1.04) | -67.50 (±1.51) | 8 | <0.0001 (****) |

| Figure | Label | Genotype | $[Ca^{2+}]_E$ (mM) | PhTx? | mEPSP amplitude (mV) | EPSP amplitude (mV) | QC | Rinput (MΩ) | Resting potential (mV) | n | P Value (significance; mEPSP, QC) |
|---|---|---|---|---|---|---|---|---|---|---|---|
| 3D | wild type | *w*$^{1118}$ | 0.4 | N | 1.09 (±0.06) | 35.18 (±2.51) | 32.27 (±1.32) | 5.13 (±1.24) | -66.34 (±2.98) | 7 | - |
| | | *w*$^{1118}$ | 0.4 | Y | 0.52 (±0.03) | 33.84 (±1.84) | 65.08 (±2.04) | 5.95 (±2.02) | -73.46 (±4.26) | 7 | <0.0001 (****), <0.0001 (****) |
| 3D | ID$^{-/+}$ | *w;+;kaiRID*$^{JM1}$*/+* | 0.4 | N | 1.12 (±0.04) | 35.89 (±2.67) | 32.04 (±1.38) | 7.61 (±0.88) | -64.33 (±1.64) | 6 | - |
| | | *w;+;kaiRID*$^{JM1}$*/+* | 0.4 | Y | 0.50 (±0.02) | 32.42 (±2.16) | 64.84 (±2.49) | 5.27 (±1.94) | -63.78 (±0.98) | 6 | <0.0001 (****), <0.0001 (****) |
| 3D | Ukar$^{-/+}$ | *w;+;+;ukar*$^{CC1}$*/+* | 0.4 | N | 1.02 (±0.06) | 31.54 (±3.18) | 30.92 (±1.80) | 6.34 (±0.92) | -64.72 (±1.21) | 6 | - |
| | | *w;+;+;ukar*$^{CC1}$*/+* | 0.4 | Y | 0.53 (±0.02) | 32.83 (±1.46) | 61.94 (±1.59) | 7.24 (±1.73) | -65.24 (±2.34) | 6 | <0.0001 (****), <0.0001 (****) |
| 3D | Ekar$^{-/+}$ | *w;+;+;ekar*$^{CC1}$*/+* | 0.4 | N | 1.08 (±0.06) | 34.63 (±2.19) | 32.06 (±1.17) | 8.31 (±1.04) | -74.33 (±2.84) | 6 | - |
| | | *w;+;+;ekar*$^{CC1}$*/+* | 0.4 | Y | 0.52 (±0.03) | 36.74 (±0.89) | 70.65 (±0.99) | 9.68 (±1.57) | -69.08 (±3.16) | 6 | <0.0001 (****), <0.0001 (****) |
| 3D | ID$^{-/+}$; Ukar$^{-/+}$ | *w;+;kaiRID*$^{JM1}$*/+; ukar*$^{CC1}$*/+* | 0.4 | N | 1.06 (±0.05) | 34.52 (±2.41) | 32.57 (±1.74) | 6.21 (±1.48) | -66.63 (±1.43) | 7 | - |
| | | *w;+;kaiRID*$^{JM1}$*/+; ukar*$^{CC1}$*/+* | 0.4 | Y | 0.51 (±0.04) | 17.26 (±1.48) | 33.84 (±1.31) | 7.63 (±1.32) | -68.53 (±3.15) | 7 | <0.0001 (****), 0.8125 (ns) |
| 3D | ID$^{-/+}$; Ekar$^{-/+}$ | *w;+;kaiRID*$^{JM1}$*/+; ekar*$^{CC1}$*/+* | 0.4 | N | 1.08 (±0.06) | 33.67 (±1.72) | 31.18 (±1.68) | 7.66 (±0.93) | -63.35 (±2.20) | 8 | - |
| | | *w;+;kaiRID*$^{JM1}$*/+; ekar*$^{CC1}$*/+* | 0.4 | Y | 0.52 (±0.02) | 30.49 (±0.76) | 58.63 (±0.84) | 9.38 (±1.37) | -66.64 (±1.06) | 9 | <0.0001 (****), <0.0001 (****) |

| Figure | Label | Genotype | $[Ca^{2+}]_E$ (mM) | GluRIIA$^{-/-}$? | mEPSP amplitude (mV) | EPSP amplitude (mV) | QC | Rinput (MΩ) | Resting potential (mV) | n | P Value (significance: mEPSP, QC) |
|---|---|---|---|---|---|---|---|---|---|---|---|
| 3F | wild type | *w*$^{1118}$ | 0.4 | N | 1.11 (0.02) | 35.26 (0.61) | 31.77 (0.51) | 7.69 (±0.82) | -72.73 (±3.25) | 7 | - |
| | | *w;GluRIIA*$^{PV3}$ | 0.4 | Y | 0.46 (0.04) | 32.15 (3.48) | 69.89 (4.37) | 9.13 (±0.56) | -73.94 (±1.35) | 7 | <0.0001 (****), <0.0001 (****) |
| 3F | ID$^{-/+}$ | *w;+;kaiRID*$^{JM1}$*/+* | 0.4 | N | 1.07 (0.03) | 33.46 (1.68) | 31.28 (0.91) | 5.81 (±0.71) | -68.56 (±0.56) | 6 | - |
| | | *w;GluRIIAPV3; kaiRID*$^{JM1}$*/+* | 0.4 | Y | 0.52 (0.02) | 31.27 (2.12) | 60.13 (2.35) | 6.96 (±0.53) | -67.90 (±2.53) | 6 | <0.0001 (****), <0.0001 (****) |
| 3F | Ukar$^{-/+}$ | *w;+;+;ukar*$^{CC1}$*/+* | 0.4 | N | 1.12 (0.05) | 30.87 (1.64) | 27.56 (0.85) | 5.73 (±0.72) | -67.27 (±3.71) | 6 | - |
| | | *w; GluRIIAPV3;+; ukar*$^{CC1}$*/+* | 0.4 | Y | 0.47 (0.04) | 29.57 (2.59) | 62.91 (3.18) | 12.91 (±1.03) | -68.83 (±2.82) | 6 | <0.0001 (****), <0.0001 (****) |
| 3F | Ekar$^{-/+}$ | *w;+;+;ekar*$^{CC1}$*/+* | 0.4 | N | 1.09 (0.04) | 32.53 (1.85) | 29.84 (0.98) | 7.93 (±0.54) | -64.69 (±1.08) | 8 | - |
| | | *w; GluRIIA*$^{PV3}$*;+; ekar*$^{CC1}$*/+* | 0.4 | Y | 0.51 (0.03) | 36.48 (0.43) | 71.53 (1.06)) | 9.41 (±0.83) | -65.26 (±1.56) | 8 | <0.0001 (****), <0.0001 (****) |
| 3F | ID$^{-/+}$; Ukar$^{-/+}$ | *w;+;kaiRID*$^{JM1}$*/+; ukar*$^{CC1}$*/+* | 0.4 | N | 1.04 (0.06) | 33.19 (3.11) | 31.91 (0.49) | 7.22 (±0.98) | -63.87 (±0.62) | 8 | - |
| | | *w; GluRIIA*$^{PV3}$*; kaiRID*$^{JM1}$*/+;ukar*$^{CC1}$*/+* | 0.4 | Y | 0.53 (0.04) | 16.98 (1.25) | 32.04 (1.73) | 8.59 (±0.51) | -69.29 (±3.26) | 8 | <0.0001 (****), 0.2235 (ns) |
| 3F | ID$^{-/+}$; Ekar$^{-/+}$ | *w;+;kaiRID*$^{JM1}$*/+; ekar*$^{CC1}$*/+* | 0.4 | N | 1.05 (0.02) | 35.61 (1.93) | 33.91 (1.36) | 9.16 (±0.43) | -66.19 (±2.51) | 10 | - |
| | | *w; GluRIIA*$^{PV3}$*; kaiRID*$^{JM1}$*/+;ekar*$^{CC1}$*/+* | 0.4 | Y | 0.45 (0.03) | 15.51 (1.41) | 34.47 (1.81) | 11.37 (±0.64) | -65.38 (±1.11) | 10 | <0.0001 (****), 0.7276 (ns) |

| Figure | Label | Genotype | Normalized BRP puncta intensity (%-IIA) | Normalized RBP puncta intensity (%-IIA) | n | P Value (significance: BRP, RBP intensity) |
|---|---|---|---|---|---|---|
| 4 A,B | wild type | *w*$^{1118}$ | 100.00 (±9.59) | 100.00 (±8.21) | 13 | - |
| 4 A,B | wild type +GluRIIA$^{-/-}$ | *w;GluRIIA*$^{PV3}$ | 177.38 (±8.25) | 167.77 (±7.91) | 13 | <0.0001 (****), <0.0001 (****) |
| 4 A,B | Ekar$^{-/-}$ | *w;+;+;ekar*$^{CC1}$ | 100.00 (±9.69) | 100.00 (±11.86) | 11 | - |
| 4 A,B | Ekar$^{-/-}$ +GluRIIA$^{-/-}$ | *w;GluRIIA*$^{PV3}$*;+;ekar*$^{CC1}$ | 176.45 (±9.71) | 198.53 (±14.13) | 11 | <0.0001 (****), <0.0001 (****) |

| Figure | Label | Genotype | $[Ca2+]_E$ (mM) | ΔR/R | n | P Value (significance: ΔR/R) |
|---|---|---|---|---|---|---|
| 4 C,D | wild type | *w;OK319/+;UAS-Syt::mScarlet3::GCaMP8m/+* | 0.8 | 0.378 (±0.025) | 12 | - |
| 4 C,D | wild type +GluRIIA$^{-/-}$ | *w;OK319,IIA*$^{PV3}$*/IIA*$^{PV3}$*;UAS-Syt::mScarlet3::GCaMP8m/+* | 0.8 | 0.603 (±0.047) | 18 | <0.0001 (****) |
| 4 C,D | Pre > Ekar$^{RNAi}$ | *w;OK319;UAS-Ekar*$^{RNAi}$*/ UAS-Syt::mScarlet3::GCaMP8m* | 0.8 | 0.342 (±0.031) | 12 | - |

| | | | | | | |
|---|---|---|---|---|---|---|
| 4 C,D | Pre > Ekar$^{RNAi}$ +GluRIIA$^{-/-}$ | *w;OK319,IIA$^{PV3}$;UAS-Ekar$^{RNAi}$/ UAS-Syt::mScarlet3::GCaMP8m* | 0.8 | 0.389 (±0.028) | 17 | 0.7936 (ns) |

| Figure | Label | Genotype | [Ca$^{2+}$]$_E$ (mM) | mEPSP amplitude (mV) | EPSP amplitude (mV) | QC | Rinput (MΩ) | Resting potential (mV) | n | P Value (significance: mEPSP, QC) |
|---|---|---|---|---|---|---|---|---|---|---|
| 4 Supplemental | Pre > Ekar$^{RNAi}$ | *w;OK319/+; UAS-Ekar$^{RNAi}$/+* | 0.4 | 1.06 (±0.03) | 32.98 (±1.91) | 31.11 (±1.10) | 9.36 (±0.72) | -68.51 (±1.09) | 7 | - |
| 4 Supplemental | Pre > Ekar$^{RNAi}$ +GluRIIA$^{-/-}$ | *w;OK319,IIA$^{PV3}$/IIA$^{PV3}$; UAS-Ekar$^{RNAi}$/+* | 0.4 | 0.52 (±0.02) | 17.39 (±1.27) | 33.44 (±0.73) | 7.21 (±0.81) | -67.19 (±2.63) | 13 | <0.0001 (****), 0.8645 (ns) |
| 4 Supplemental | Post > Ekar$^{RNAi}$ | *w;G14/+; UAS-Ekar$^{RNAi}$/+* | 0.4 | 1.04 (±0.03) | 30.36 (±2.72) | 29.19 (±1.57) | 10.05 (±1.27) | -69.30 (±1.52) | 6 | - |
| 4 Supplemental | Post > Ekar$^{RNAi}$ +GluRIIA$^{-/-}$ | *w;G14,IIA$^{PV3}$/IIA$^{PV3}$; UAS-Ekar$^{RNAi}$/+* | 0.4 | 0.49 (±0.03) | 31.54 (±1.16) | 64.37 (±0.67) | 8.36 (±0.83) | -65.70 (±1.68) | 6 | <0.0001 (****), <0.0001 (****) |

### KEY RESOURCES TABLE

| REAGENT/RESOURCE | | SOURCE | IDENTIFIER |
|---|---|---|---|
| **Chemicals, peptides, and recombinant proteins** | | | |
| Philanthotoxin-433 | | Sigma-Aldrich | 276684-27-6 |
| **Antibodies** | **Dilution** | | |
| Mouse anti-BRP (nc82) | 1:500 | Developmental Studies Hybridoma Bank (DSHB) | AB_2314866 |
| Mouse anti-GFP (4C9) | 1:200 | DSHB | AB_2617422 |
| Guinea pig anti-RBP | 1:2000 | (He et al., 2023) | |
| Guinea pig anti-vGlut | 1:2000 | (Chen et al., 2017) | |
| Alexa Fluor 647 conjugated Goat anti-Horseradish Peroxidase | 1:400 | Jackson ImmunoReserach Laboratories (Jackson) | 123-605-021 |
| Cy3-conjugated secondary antibodies | 1:400 | Jackson | 106-165-003 |
| Alexa Fluor 488 conjugated secondary antibodies | 1:400 | Jackson | 715-545-150 |
| Phalloidin-Alexa Fluor 647 | 1:1000 | Thermo Fisher | A22287 |
| **Drosophila Strains** | | | |
| *w*$^{1118}$ | | BDSC | 5905 |
| *GluRIIA*$^{PV3}$ | | (Han et al., 2023) | |
| *GluRIIA-T2A-Gal4* | | (Kondo et al., 2020) | |
| *GluRIIB-T2A-Gal4* | | (Kondo et al., 2020) | |
| *GluRIIC-T2A-Gal4* | | (Kondo et al., 2020) | |
| *GluRIID-T2A-Gal4* | | (Kondo et al., 2020) | |
| *GluRIIE-T2A-Gal4* | | (Kondo et al., 2020) | |
| *ekar*$^{CC1}$ | | This study | |
| *ekar*$^{CC2}$ | | This study | |
| *ekar-T2A-Gal4* | | This study | |
| *ekar-Gal4* | | BDSC | 22661 |
| *ukar*$^{CC1}$ | | This study | |
| *ukar*$^{CC2}$ | | This study | |
| *ukar*$^{CRIMIC}$ (Gal4) | | BDSC | 97180 |
| *kaiRID*$^{2}$ | | BDSC | 22962 |
| *kaiRID*$^{JM1}$ | | This study | |
| *kaiRID*$^{CRIMIC}$ (Gal4) | | BDSC | 98475 |
| *grik*$^{JM1}$ | | This study | |
| *grik*$^{JM2}$ | | This study | |
| *grik-Gal4* | | BDSC | 24332 |
| *clumsy*$^{NT1}$ | | This study | |
| *clumsy*$^{NT2}$ | | This study | |
| *clumsy*$^{CRIMIC}$ (Gal4) | | BDSC | 606523 |
| *gluRIA*$^{NT1}$ | | This study | |
| *gluRIA*$^{NT2}$ | | This study | |
| *gluRIA-T2A-Gal4* | | BDSC | 84640 |
| *gluRIB*$^{NT1}$ | | This study | |
| *gluRIB*$^{NT2}$ | | This study | |
| *gluRIB-Trojan-Gal4* | | BDSC | 76135 |
| *NMDAR1*$^{XC1}$ | | (Li et al., 2021) | |
| *NMDAR1*$^{XC3}$ | | (Li et al., 2021) | |
| *NMDAR1-T2A-Gal4* | | BDSC | 84669 |
| *NMDAR2*$^{NT1}$ | | This study | |
| *NMDAR2*$^{NT2}$ | | This study | |
| *NMDAR2-T2A-Gal4* | | BDSC | 84670 |

| | | |
|---|---|---|
| *GluClα¹* | BDSC | 38025 |
| *GluClα³* | BDSC | 6353 |
| *GluClα Ile27 to Val mutant* | (Zak et al., 2024) | |
| *GluClα-Trojan-Gal4* | BDSC | 77841 |
| *mGluRA^NT1* | This study | |
| *mGluRA^NT2* | This study | |
| *mGluRA-Trojan-Gal4* | BDSC | 77721 |
| *OK319-Gal4* | (Sweeney et al., 1995) | |
| *G14-Gal4* | (Aberle et al., 2002) | |
| *UAS-mCD8::tdGFP* | BDSC | 32194 |
| *UAS-Syt::mScarlet3::GCaMP8m* (Scar8m) | (Chen et al., 2026) | |
| *bam-cas9* | BDSC | 97543 |
| *nos-cas9* | BDSC | 78782 |
| **Software and Algorithms** | | |
| Clampex | Molecular Devices | 10.7 |
| Mini Analysis | Synaptosoft | 6.0.7 |
| Axon pClamp Clampfit | Molecular Devices | 10.7 |
| NIS-Elements Software | Nikon Instruments | 5.41.02 |
| Huygens Essential | Scientific Volume Imaging | 25.04 |
| Jupyter Notebook | Anaconda | 6.0.1 |
| Python | https://www.python.org/ | |
| Cafire | (Chen et al., 2026) | |
| Graphpad Prism | Graphpad | 10.0.1 |
| Excel | Microsoft | 2022 |
